# ICER controls HIV-1 infection and replication in elite controllers

**DOI:** 10.1101/2021.06.30.450567

**Authors:** Zhenwu Luo, Min Li, Taiwei Li, Zongyang Lv, Zhiwei Ye, William J. Cisneros, Jie Zhang, Lingmin Yuan, Judd F. Hultquist, Stephen A Migueles, Jian Zhu, Wei Jiang

**Affiliations:** Department of Microbiology and Immunology, Medical University of South Carolina, Charleston, SC, USA; Department of Pathology, Ohio State University College of Medicine, Columbus, OH, USA; Department of Biochemistry & Molecular Biology, Hollings Cancer Center, Medical University of South Carolina, Charleston, SC, USA; Department of Biochemistry & Structural Biology, University of Texas Health Science Center at San Antonio, San Antonio, TX, USA; Department of Cell and Molecular Pharmacology and Experimental Therapeutics, Medical University of South Carolina, Charleston, SC, USA; Division of Infectious Diseases, Northwestern University Feinberg School of Medicine, Chicago, IL, USA; Laboratory of Immunoregulation, National Institute of Allergy and Infectious Diseases, National Institutes of Health, Bethesda, MD, USA; Division of Infectious Diseases, Department of Medicine, Medical University of South Carolina, Charleston, SC, USA

**Keywords:** HIV, elite controllers, CREM/ICER gene, host factor, alternative RNA splicing

## Abstract

A rare subset of HIV-infected individuals, termed elite controllers (ECs), can maintain long-term control over HIV replication in the absence of antiretroviral therapy (ART). To elucidate the biological mechanism of resistance to HIV replication at the molecular and cellular levels, we performed RNA sequencing and identified alternative splicing variants from ECs, HIV-infected individuals undergoing ART, ART-naïve HIV-infected individuals, and healthy controls. Differential gene expression patterns that are specific to ECs and may influence HIV resistance were identified, including alternative RNA splicing and exon usage variants of the CREM/ICER gene (cAMP-responsive element modulator/inducible cAMP early repressors). The knockout and knockdown of specific ICER exons that were found to be upregulated in ECs resulted in significantly increased HIV infection in CD4+ T cell line and primary CD4+ T cells. Overexpression of ICER isoforms decreased HIV infection in primary CD4+ T cells. Furthermore, ICER regulated HIV-1 LTR promoter activity in a Tat-dependent manner. Together, these results suggest that ICER is an HIV host factor that may contribute to HIV resistance of ECs. These findings will help elucidate the mechanisms of HIV control by ECs and may yield a new approach for treatment of HIV.

## INTRODUCTION

A small group of HIV-1-infected individuals, termed elite controllers (ECs), display control of HIV replication in the absence of antiretroviral therapy (ART) [1]. This is in contrast to most humans, in whom HIV infection progresses to acquired immunodeficiency syndrome (AIDS) unless treated with ART. ECs can spontaneously control HIV-1 replication without ART and maintain HIV RNA levels in the blood below the level of detection [1, 2]. Some aspects of the natural course of infection in ECs that contribute to this resistance have been described, including protective HLA alleles and effective CD8+ T cell cytotoxicity, both of which inhibit viral replication [3–6]. It has also been suggested that infection with a virus that has low replicative fitness may explain viral control in some ECs, but viruses isolated from a subset of ECs have been found to be fully competent and replicate vigorously *in vitro* [7].

Together, these data suggest that the restricted replication capacity of HIV in ECs is likely driven by host-specific differences that prevent viral replication and/or disease progression. Several host restriction factors have been identified to inhibit viral replication. For example, TRIM5α [8], SAMHD1 [9], and APOBEC3G [10] were found to protect human and nonhuman primates against retroviral infection. Host-specific differences in these proteins have been linked to protection against infection and are thought to provide a significant barrier against cross-species transmission, but these host factors have not been reported to account for HIV resistance in ECs. Host factor driven mechanisms of resistance to HIV in ECs remain largely unexplored.

Previous studies found that both monocyte-derived macrophages and CD4+ T cells from the HIV controllers had low susceptibility to HIV infection *in vitro* [11, 12]. However, the restriction of viral replication in CD4+ T cells from HIV controllers can be overcome by high-dose viral challenges [11, 13]. Additionally, some reports have found that CD4+ T cells from ECs can be infected by autologous or laboratory strains of HIV-1 [13, 14]. A previous study looking to identify potential host factors regulating viral replication compared with the transcriptomes of CD4+ T cells between ECs and ART-naïve HIV-infected individuals [15]. While transcriptional profile broadly correlated with viral set point of the patients, it was unclear if this profile was driven by different levels of infected T cells being sampled or by systemic difference within each individual. Compared to CD4+ T cells, myeloid-derived cells, such as monocytes, macrophages, and dendritic cells, are generally more resistant to HIV-1 infection, in part due to the increased expression of innate immune factors [16].

Based on these previous studies, we hypothesized that specific genetic programs in monocytes from ECs might provide information on cell resistance to infection. As monocytes are not the main target of infection, sequencing of the monocyte population would further avoid confounders driven by infection-dependent changes. To test this hypothesis, we performed transcriptomics and analyzed gene expression profiles of monocytes derived from ECs, HIV-infected individuals undergoing ART, ART-naïve HIV-infected individuals, and healthy controls. We focused on the genes that exhibited increased or decreased expression specifically in ECs relative to the other three control groups (non-ECs). Here, we show increased expression of specific exons of the ICER (cAMP-responsive element modulator/inducible cAMP early repressors) locus specifically in ECs. Knockout of ICER in CD4+ T cell line increased HIV infection; overexpression or knockdown of these isoforms in primary CD4+ T cells decreased and increased HIV infection, respectively. These findings suggest that specific genetic programs in ECs, including altered splicing of ICER, may regulate cellular resistance to HIV infection.

## RESULTS

### Screening of the genes controlling HIV infection in ECs

To identify host factors specific to ECs that might be involved in controlling HIV infection, we performed RNA sequencing (RNA-Seq) to compare the transcriptome of monocytes from ECs and non-ECs. Peripheral blood mononuclear cells were obtained from 44 individuals (**Supplementary Table 1**), comprised of 10 ECs, 12 aviremic HIV-infected ART-treated individuals, 11 HIV-infected ART-naïve individuals, and 11 healthy controls without HIV infection. Monocytes were enriched from each sample, and RNA was extracted for RNA-Seq. To explore critical molecular factors and pathways, we compared distinct gene expression in ECs with those in the other three control groups (non-ECs) (**Fig. 1A**). Overall, 2836 genes exhibited significantly increased or decreased expression in ECs (log2 fold change > 1.4, q-value < 0.05) compared to ART-treated individuals, ART-naïve individuals, or healthy control individuals. Of these, we found 53 candidate genes that were differentially expressed in ECs compared to each non-EC group (**Supplementary Table 2**) that might contribute to the different outcomes of HIV infection in ECs independent of HIV infection or ART-status.

**Figure 1.**
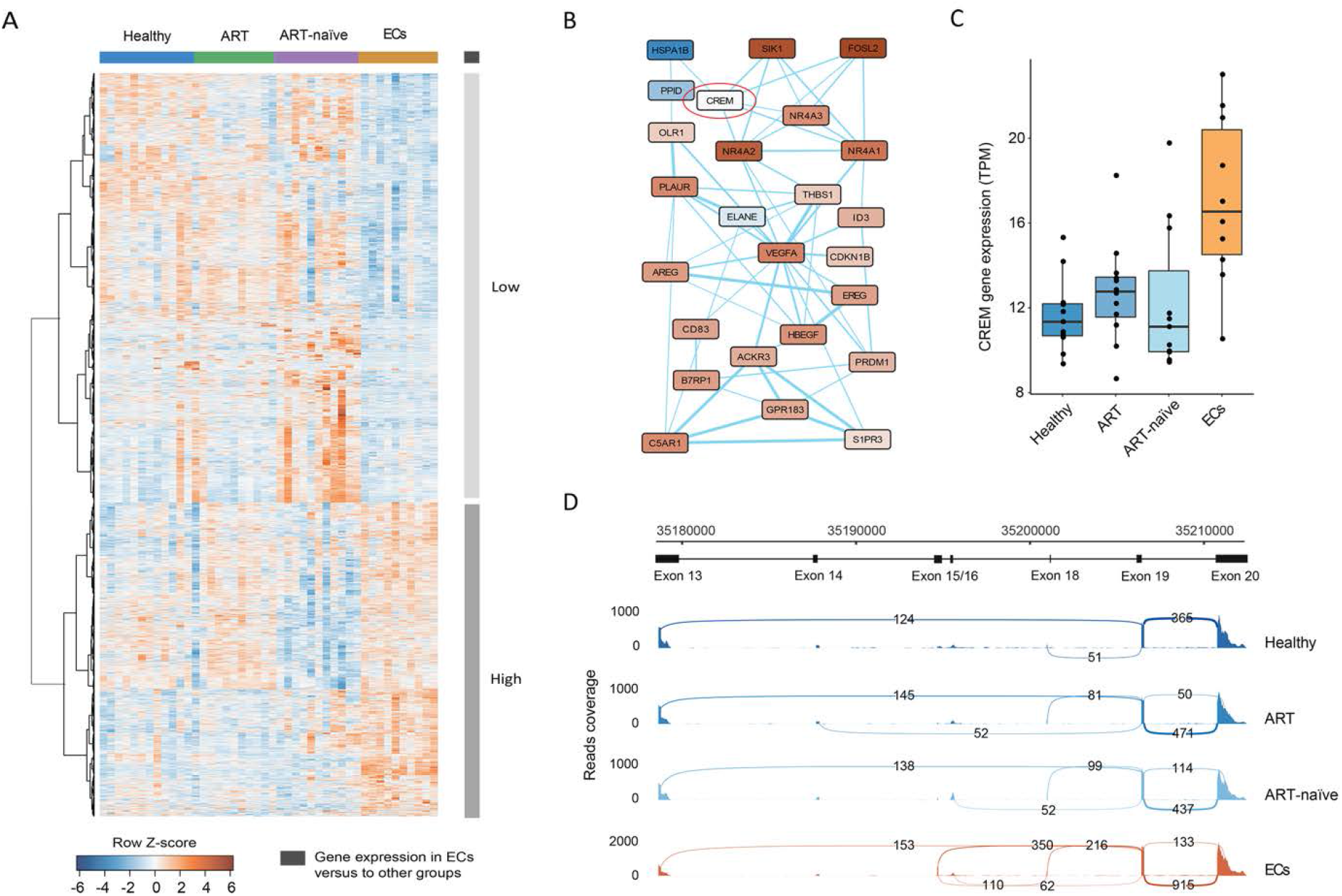
Unique gene expression in ECs. Gene expression was analyzed by RNA-seq in purified monocytes from ECs, ART-treated and viral-suppressed (ART), and ART-naïve individuals, as well as healthy controls (Healthy). (A) Heatmap demonstrating hierarchical clustering of the distinctly expressed genes of transcripts per million (TPM) in the four study groups. In total, 2836 genes met the gene selection criteria (log2-fold change > 1.4, FDR adjusted P < 0.05), which were significantly different from at least one of the ECs/ART, ECs/ART-naive, and ECs/Healthy comparisons. (B) Functional gene network showing the genes increased (red) or decreased (blue) in ECs compared with the other three groups simultaneously. The edge width reflects the interaction score between two genes. The minimum required interaction score was > 0.3. (C) Overall RNA expression level (TPM) of CREM in ECs, ART, ART-naïve, and healthy controls. (D) Sashimi plot of CREM exons 13-20, displaying raw mapped RNA sequencing reads. Each group contained ten pooled samples.

Next, the identified genes were subjected to gene function and network linkage analyses. Using Gene Ontology (GO) to functionally characterize these 53 candidate genes, we found that “cell cycle”, “cellular process and activation”, and “apoptotic process” represented the top functional entities that were differentially expressed and contained most of the genes distinguishing ECs from non-ECs (**Fig. S1A**). Using STRING to analyze interaction networks, we identified a notable cluster that includes several of the most markedly variable genes, such as the most highly upregulated genes SKI1B and NR4A2, and the most highly downregulated gene HSPA1B **(Fig. 1B)**. To determine if the differentially expressed genes in ECs confer HIV resistance phenotypes to other cells, we overexpressed the 12 most significantly altered genes (**Supplementary Table 3**). Each of the genes was cloned into a lentiviral expression vector and used to make high titer lentivirus. Interferon-alpha 1 (IFNA1) was used as an HIV-resistant positive control [17, 18]; CD8 or an empty vector were used as negative controls. HuT78 cells were transduced and cultured for three weeks for stable expression of the transduced genes prior to challenge with HIV-1 NL4-3 at the multiplicity of infections (MOIs). Percent infected cells were quantified by intracellular HIV p24 staining 72 h after HIV infection. Except for IFNA1, which showed significant resistance to HIV challenge, we failed to observe differences in HIV infection after overexpression of the differentially expressed genes **(Fig. S1B)**.

In the gene interaction network cluster, the CREM gene interacted with both the most highly upregulated and downregulated genes. While not among the most highly differentially expressed genes, the overall RNA expression level of CREM was significantly increased in ECs compared to that in non-ECs (**Fig. 1C**). CREM encodes the cAMP-responsive element modulator, a bZIP transcription factor that plays a critical role in cAMP-regulated signal transduction. CREM has a large number of alternatively spliced transcript variants that are regulated posttranscriptionally (> 40 in humans). Some of these isoforms serve as transcriptional activators, and the others work as repressors [19]. Globally, to compare compositional dissimilarity among the alternative splicing events, unsupervised PCA analysis was applied to calculate sample distances. ECs had extinct alternative splicing events when compared with the other non-EC groups (**Fig. S1C**). Indeed, visualizing the RNA sequencing data from ECs in Integrative Genomics Viewer (IGV) showed several unique splicing junction events, particularly on chr10:35,195,000-35,213,000, which spanned the CREM gene from exon 15 to exon 20 (**Fig. 1D**). Based on the central nature of CREM in the regulatory network and its known function as a transcriptional activator/repressor, we hypothesized that CREM might be the key upstream regulatory molecule that regulates differentially expressed genes in ECs.

### CREM/ICER isoform enriched variants and alternative splicing in ECs

The human CREM gene contains 20 exons, which use different promoters and transcription factors to generate complex alternative messenger RNA (mRNA) splicing events (**Fig. S1E**). The alternatively spliced variants generate different CREM isoforms that may exert opposing effects on target gene expression depending on the absence or presence of the transactivating domains [20]. Therefore, identification of the exon usage and alternative splicing isoforms is essential to interpreting changes in RNA expression from this locus. To determine which transcripts may be differentially regulated, we compared unique reads to each isoform and calculated expression of each transcript per million reads (TPM). Four specific CREM isoforms were found to be enriched in ECs over non-ECs (**Fig. S1D**). Notably, all of the isoforms enriched in ECs were inducible cAMP early repressors or ICERs. These isoforms displayed preferential usage of exons 15 and 16 (**Fig. 2A**). This same enrichment was observed when calculated as isoform fraction (**Fig. 2B**). On the other hand, two CREM isoforms were enriched in non-ECs compared to ECs, one of them encoding an IncRNA and the other spanning exon 2 to exon 20 (**Fig. 2A**). We also quantified the alternative splicing events within CREM by analyzing the percent spliced in (PSI or Ψ, **Fig. 2C**). Consistent with the IF analysis, the PSI results showed that ECs had an alternative first exon (AFE) preference for usage of exon 16 (median PSI = 90.2%), short exon 16 (median PSI = 89.0%), and exon 15 (median PSI = 4.6%). In ECs, when the first exon of CREM was exon 15, exon 18 had a highly skipped exon (SE) fraction (median PSI = 25.2%).

**Figure 2.**
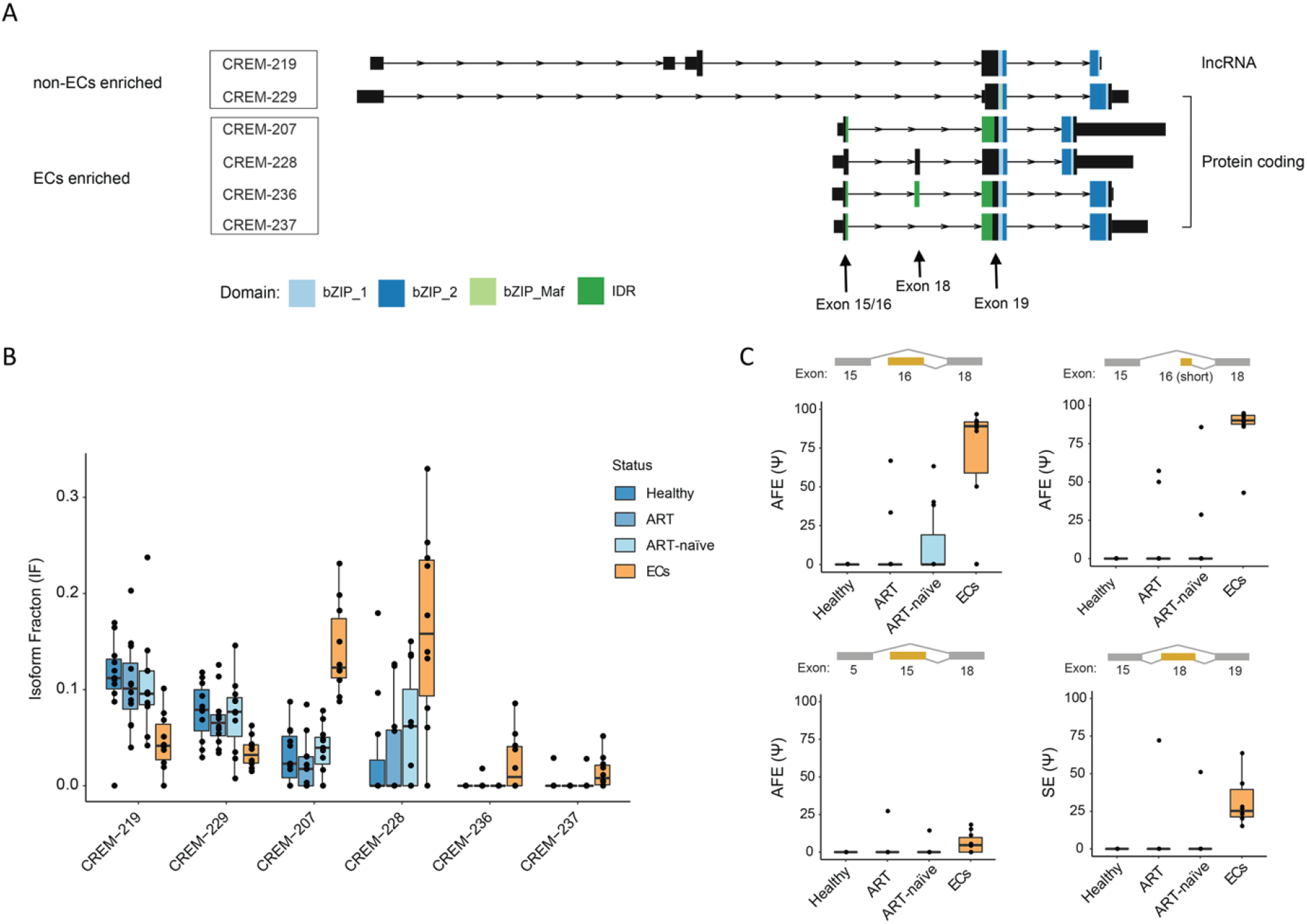
EC-specific CREM/ICER isoforms and alternative splicing based on RNA sequencing. (A) Schematic illustration of CREM/ICER displaying the six relevant transcript isoforms (Ensembl ID) with their predicted functional domain structures. The expression of these six CREM/ICER isoforms was significantly increased or decreased in ECs compared to all three control groups (non-ECs). (B) The frequencies of six CREM/ICER transcript isoform fractions (IF) among all CREM/ICER isoforms in the four study groups. (C) Splicing patterns within the EC-relevant CREM/ICER genes quantified by percent spliced in (PSI or Ψ) values in the four study groups; the PSI with a significant difference is shown. The preference alternative first exon (AFE) usage and skipped exon (SE) in ECs are highlighted.

### Knockout of ICER exons using CRISPR-Cas9 increase HIV infection in CD4+ T cell line

Given the many CREM/ICER isoforms, we used a CRISPR-Cas9 system to knock out the CREM/ICER gene in HuT78 cells to explore its role in HIV replication. We designed several synthetic single-guide RNAs (sgRNAs) against ICER exons 15 and 16 **(Supplementary Table 4).** Exon 18 and 19, which were commonly used in CREM/ICER isoforms (**Fig. S1E**), were also knocked out. Exon 13, which was used in most CREM isoforms but absent in ICER isoforms (**Fig. S1E**), was chosen as control. ICOSLG, one of the most upregulated genes in ECs **(Fig. 1B)**, didn’t change the HIV infection after overexpression in HuT78 cells (**Fig. S1B**), was chosen as a negative control. Individual CREM/ICER exon or ICOSLG gene knockout clones were obtained through single-cell sorting and screened by sequencing. We selected three clones for each exon or gene knockout. Next, the cell clones were challenged with HIV NL4-3, and infection was quantified by intracellular HIV p24 staining and flow cytometry at 72 h. The cell clones carrying the frameshift in ICER were more vulnerable to HIV challenge (**Fig. 3A**). To confirm these results, representative mutant clones that had frameshifts in each exon were challenged with increasing doses of HIV-1 NL4-3. The productive infection was quantified. As observed above, each ICER exon’s knockout significantly promoted infection on day 3 (**Fig. 3B, 3C**). Similar results were obtained using the HIV NL4-3 virus to challenge cells with each frameshift mutation after cultivation for 6 days (**Fig. 3B, 3C**). The cell clones suffered significantly decreasing cell counts due to HIV-related cell death by the day 6 and day 9 timepoint (**Fig. 3B, 3D**).

**Figure 3.**
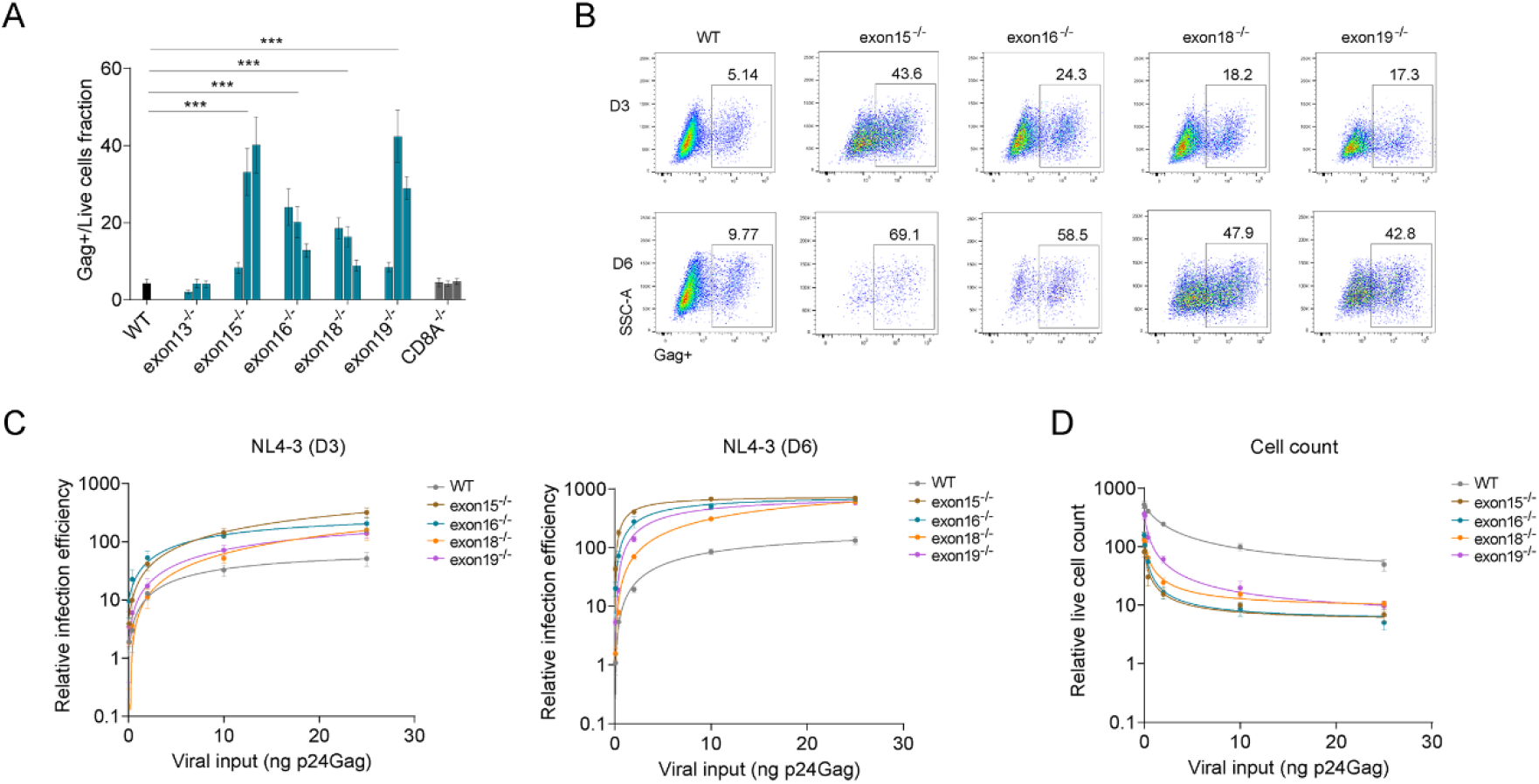
Knockout of ICERs increase HIV infection. CREM/ICER exon 13, 15, 16, 18, or 19 in HuT78 cells were knockout using the CRISPR/Cas9 system. The viral replication efficiency on each CREM/ICER exon knockout HuT78 cell clones were evaluated by HIV gag staining using flow cytometry. (A) The fold changes of HIV-1 infection in every three knockout clones on CREM/ICER exon 13, 15, 16, 18, or 19 or CD8A, compared to wild-type HuT78 cells on day 3 after HIV NL4-3 infection. (B) Dot plots representing Gag+ cells on day 3 and day 6 after challenging HIV NL4-3 with individual exon knockout in the HuT78 cell line. The knockout on ICER exons 15, 16, 18, or 19 in HuT78 cells generated different resistance patterns to HIV infection. Mean levels of five replicates from three independent experiments are shown. (C, D) The representative cells with the knockout of ICER exon 15, 16, 18, or 19 were infected with increasing viral inputs of HIV NL4-3. The infection efficiency was monitored on day 3 (C) and day 6 by measuring Gag+ cells. The live cell counts were measured on day 9 (D). Mean ± SD. Five replicates from three independent experiments. One-way ANOVA followed by Holm-Sidak’s multiple comparisons test. \*\*\**p* < 0.001.

### Knockdown of ICER exons using shRNA leads to increased HIV infection in primary CD4+ T cells

To determine the contribution of ICER to HIV infection in primary CD4+ T cells, short hairpin RNA (shRNA)-mediated silencing was used to suppress ICER expression (**Supplementary Table 5**). Lentiviruses containing shRNA that target distinct exons were used to transduce primary CD4+ T cells isolated from 4 different non-HIV-infected donors. The efficiency and specificity of ICER were assessed by immunoblotting (**Fig. 4A**). The shRNA against exons 15 and 19 had high knockdown efficiency of ICER, while shRNA against exons 16 and 18 had low knockdown efficiency similar to the non-targeting control (**Fig. 4A**). Similar levels of cell viability were observed in CD4+ T cells after ICER knockdown compared to control shRNA-transduced CD4+ T cells (**Fig. S2A**). Transduced cells were subsequently infected with HIV-1 AD8 (CCR5-tropic) or HIV-1 NL4-3 (CXCR4-tropic) virus, and infected cells were quantified by intracellular HIV p24 staining on days 3 and 5 post-infection. Upon ICER depletion, both HIV-1 AD8 and NL4-3 infection were dramatically increased in primary T cells from all four donors compared to the cells that were transduced with control shRNA (**Fig. 4B-4D**). The three shRNAs with the highest knockdown efficiencies, shRNA 15-1, shRNA 15-2, and shRNA 19-2, had an average of 5.0, 9.1, and 5.8-fold increases in HIV-1 AD8 infection, respectively, compared with control shRNA on day 3 after viral challenge (**Fig. 4C**). Furthermore, shRNA 15-1 and shRNA 19-2 had an average of 4.6 and 2.8-fold increases in HIV NL4-3 infection, respectively, compared with control shRNA on day 3 post-infection (**Fig. 4D**). Similar results were obtained using the HIV AD8 and NL4-3 virus to challenge cells with each knockdown after cultivation for five days (**Fig. S2B-S2C**).

**Figure 4.**
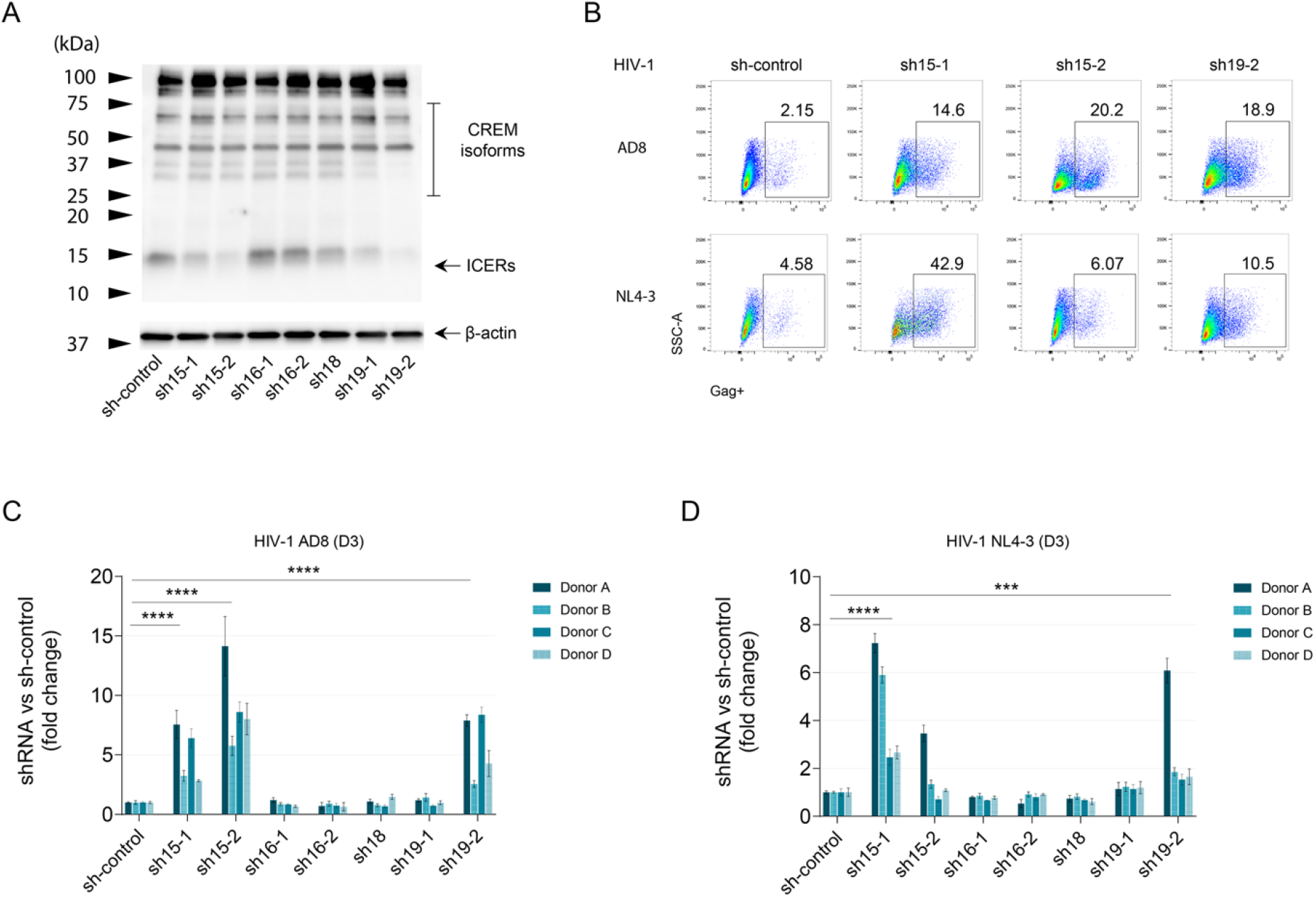
Increased HIV infection in primary CD4+ T cells upon ICER knockdown. Primary CD4+ T cells were activated using anti-CD2/CD3/CD28, transduced with ICER shRNAs against exons 15 (sh15-1, sh15-2), 16 (sh16-1, sh16-2), 18 (sh18), or 19 (sh19-1, sh19-2). The empty lentivirus was used as a control (sh-control). The transduced cells were selected using puromycin. (A) The expression of ICER after shRNA knockdown in primary CD4+ T cells. (B) Dot plots representing percents of Gag+ cells on day 3 after HIV AD8 or HIV NL4-3 infection upon ICER exon 15 or 19 knockdown in primary CD4 T cells. (C, D) Summarized results of HIV AD8 (C) or NL4-3 (D) infection after shRNA knockdown of ICER exon 15, 16, 18, or 19 in primary CD4+ T cells isolated from four different donors. Results are shown as the fold change of percent infection in ICER shRNA transduced cells compared to those of sh-control transduced cells on day 3 after viral challenges. Each histogram bar represents mean ± SD of triplicates for each donor. One-way ANOVA followed by Holm-Sidak’s multiple comparisons test. *** *p* < 0.001, \*\*\*\**p* < 0.0001.

To further verify the effect of ICER exon knockdown in inhibiting HIV infection, we examined a range of different HIV-1 strains. Human primary CD4+ T cells were challenged with different strains of HIV-1 after silencing ICER expression. The tested HIV included CCR5-tropic (JR-CSF and BaL), CXCR4/CCR5-tropic (89.6), and CXCR4-tropic (IIIB) HIV-1 strains. Treatment with shRNA 15-1 and 15-2 (knockdown of exon 15) and shRNA 19-2 (knockdown of exon 19) increased HIV infection of all tested strains in primary CD4+ T cells from multiple independent donors on day 3 post-infection (**Fig. 5A-5D**) and day 5 post-infection (**Fig. S3A-S3D**).

**Figure 5.**
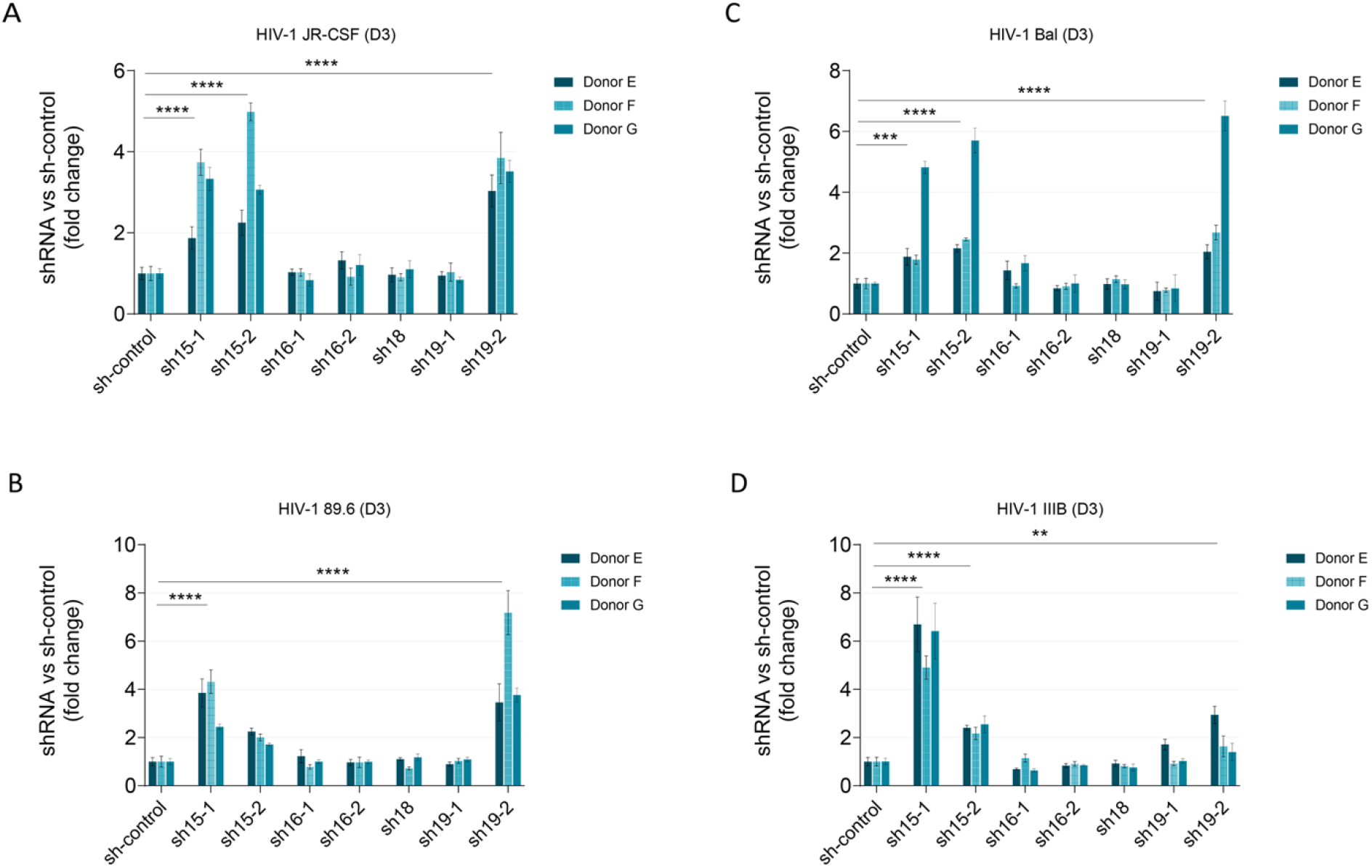
ICER controls infections with different strains of HIV-1. Primary CD4+ T cells were activated using anti-CD2/CD3/CD28, transduced with ICER shRNAs against exons 15 (sh15-1, sh15-2), 16 (sh16-1, sh16-2), 18 (sh18), or 19 (sh19-1, sh19-2), and selected by puromycin; empty lentivirus was used as a control (sh-control). The transduced cells were then infected with HIV JR-CSF (A), BaL (B), 89.6 (C), and IIIB (D). The viral replication efficiency was evaluated by p24 staining on day 3 after HIV infection. Results are displayed as the fold changes of percent infection in ICER shRNA transduced cells compared to those of sh-control transduced cells from three different donors. Each bar represents mean ± SD of triplicates for each donor. One-way ANOVA followed by Holm-Sidak’s multiple comparisons test. \*\**p* < 0.01, \*\*\**p* < 0.001, \*\*\*\**p* < 0.0001.

### Overexpression of ICER isoforms leads to resistance to HIV infection in primary CD4+ T cells

To better understand which ICER isoforms are influencing infection, we individually overexpressed eight different ICER isoforms in primary CD4+ T cells from 4 independent donors using a lentiviral expression system. Two isoforms of CREM were used as isoform controls and a CD8A vector was used as a negative control. The specificity of ICER isoform overexpression was assessed in HEK 293T cells (**Fig. S4A**), and the transduction efficiency of isoforms in primary CD4+ T cells was determined by immunoblot (**Fig. 6A**). Although HEK 293T cells highly expressed all of tested ICER isoforms, primary CD4+ T cells only highly expressed isoforms 5, 7, 8, 11, and 31. After transduction, the expression of ICER different isoforms didn’t affect the cell viability (**Fig. S4B**). Next, overexpression of ICER isoforms 5, 8, 11, and 31 resulted in 45.1%, 56.5%, 79.1%, and 79.5% average decreases in HIV AD8 infection on day 3, respectively compared with the CD8A transduced cells (**Fig. 6B, S4C**). The overexpression of ICER isoforms 8 and 11 resulted in 49.2% and 39.0% decreases in HIV NL4-3 infection on day 3 (**Fig. 6C, S4C**). Similar results were observed on day 5; the impact of 8, 11, and 31 on HIV-1 NL4-3 infection were more pronounced (**Fig. S4D-S4E**).

**Figure 6.**
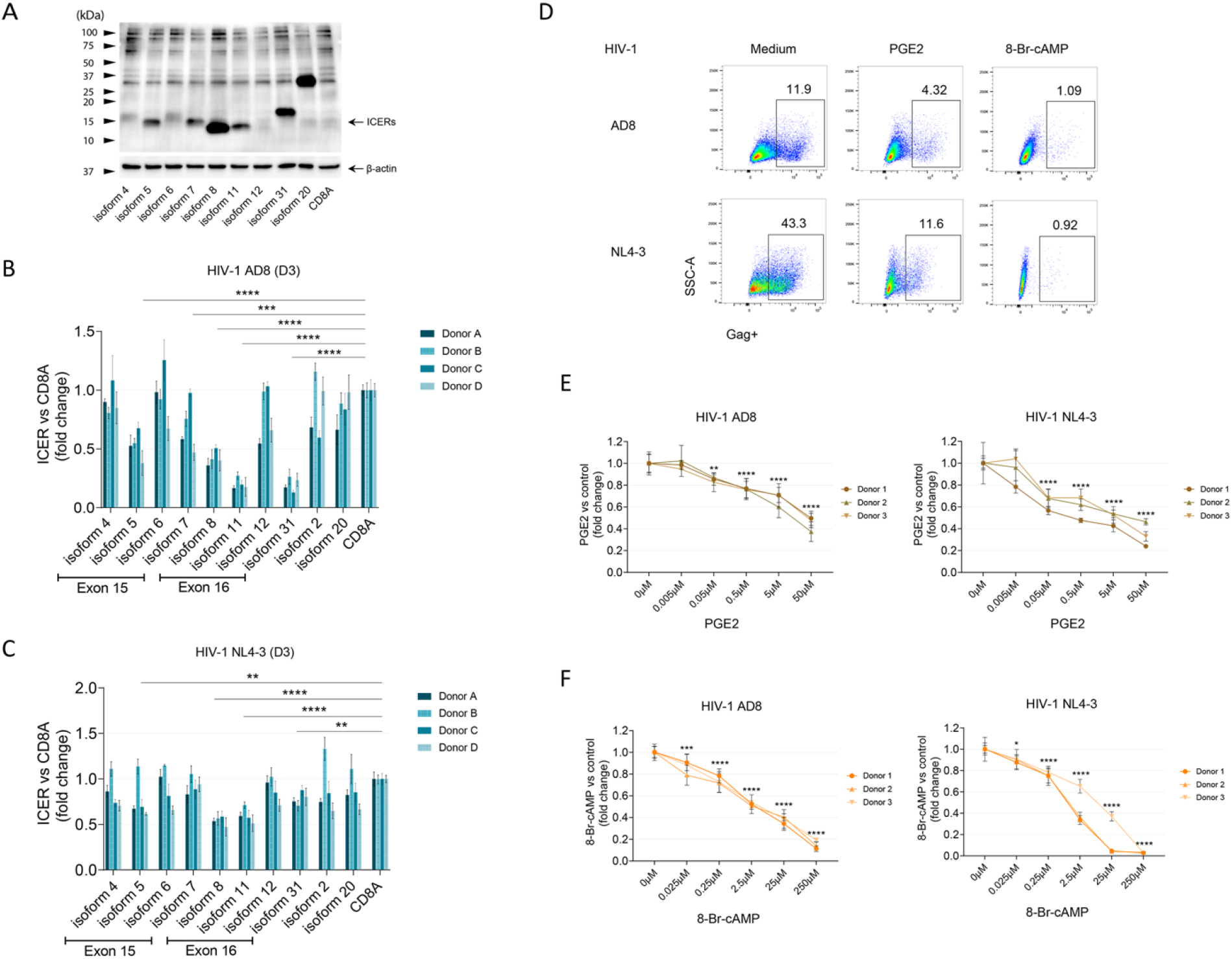
Resistance to HIV infection in human primary CD4+ T cells after ICER isoform overexpression. Primary CD4+ T cells were activated using anti-CD2/CD3/CD28, transduced with CREM/ICER isoforms with lentivirus system, and selected by puromycin. The expression of CREM/ICER isoforms using letivirus gene overexpression system was shown. (A) CREM/ICER isoform overexpression in primary CD4+ T cells. Fold changes of percent HIV AD8 (B) and NL4-3 (C) infection in human primary CD4+ T cells transduced with each CREM/ICER isoform, compared to CD8A gene transduced T cells. Histograms are shown in independent experiments performed in cells from four different donors. Each bar represents the Mean ± SD of triplicates for each donor. (D-F) Anti-CD2/CD3/CD28 activated primary CD4+ T cells were treated with PGE2 (50 μM-0.005 μM) or 8-Br-cAMP (250 μM-0.025 μM) for 16-24 h, and then infected with HIV AD8 or NL4-3 for 3 days. (D) Percent infection (% Gag+) in response to PGE2 (50 μM) or 8-Br-cAMP (250 μM) from one presentative donor. Dose-dependent effects of PGE2 (E) or 8-Br-cAMP (F) on HIV infection. Results are displayed as the fold changes of percent infection treated by ICER agonist compared to untreated cells. Histograms show results in three independent experiments performed with three different donors. Each bar represents mean ± SD of triplicates for each donor. One-way ANOVA followed by Holm-Sidak’s multiple comparisons test. * *p* < 0.05, ** *p* < 0.01, *** *p* < 0.001, **** *p* < 0.0001.

In addition to overexpression of individual ICER isoforms, the effect of small molecule agonists of ICER was assessed in the context of HIV-1 infection. Prostaglandin E2 (PGE2) and 8-Bromo-Cyclic AMP (8-Br-cAMP) have been shown to increase intracellular cAMP and induce endogenous ICER protein levels [21, 22]; they were not found to alter cell viability in the current study (**Fig. S5A-S5B**). We found both PGE2 and 8-Br-cAMP treatment increased cell resistance to HIV infection (**Fig. 6D-6F**). Treatment of human primary CD4+ T cells with PGE2 (100 μM) for 16–24 h prior to infection resulted in a 57.6% decrease in HIV AD8 infection and a 67.2% decrease in HIV NL4-3 infection, respectively, compared with untreated cells (**Fig. 6D-6F**). Notably, 8-Br-cAMP pre-treatment resulted in more dramatic inhibition of HIV infection in human primary CD4+ T cells (**Fig. 6F**), consistent with its resistance to hydrolysis by phosphodiesterases. Treatment of cells with 250 μM 8-Br-cAMP resulted in 84.8% and 97.1% decreases in HIV-1 AD8 and NL4-3 infection, respectively, compared with untreated cells (**Fig. 6F**).

### ICER regulates HIV-1 LTR promoter activity in a Tat-dependent manner

CREM/ICER has been found to regulate RNA polymerase II (RNAP II) through interactions with transcription factor IIA (TFIIA) [23]. HIV Tat was found to bind to the TAR RNA element that located in the viral 5’ Long Terminal Repeat (LTR) sequences and recruit RNAP II elongation factor to activate transcriptional elongation [24]. To evaluate the effects of CREM/ICER on RNAP II and HIV LTR promoter activity, we stably transduced TZM-B1 cells with shRNAs against CREM/ICER or over expression of CREM/ICER isoforms; IFNA1 was used as a HIV-resistant positive control. We tested the activity of HIV LTR promoter in the absence or presence of HIV Tat protein. No change of HIV LTR promoter activity was found after increasing or decreasing ICER expression (**Fig. 7A-7B**). However, in the presence of HIV-1 Tat and the depletion of ICER exon 15 led to enhanced LTR-driven luciferase reporter activity, while co-expression of HIV-1 Tat and the ICER isoforms 4, 5, and 8 decreased LTR-driven luciferase reporter activity (**Fig. 7A-7B**). These results indicated the ICER regulation of HIV-1 infection is likely through targeting HIV-1 Tat-LTR axis and viral transcription; our results are also consistent with the previous work suggesting that ICER regulates RNAP II [23].

**Figure 7.**
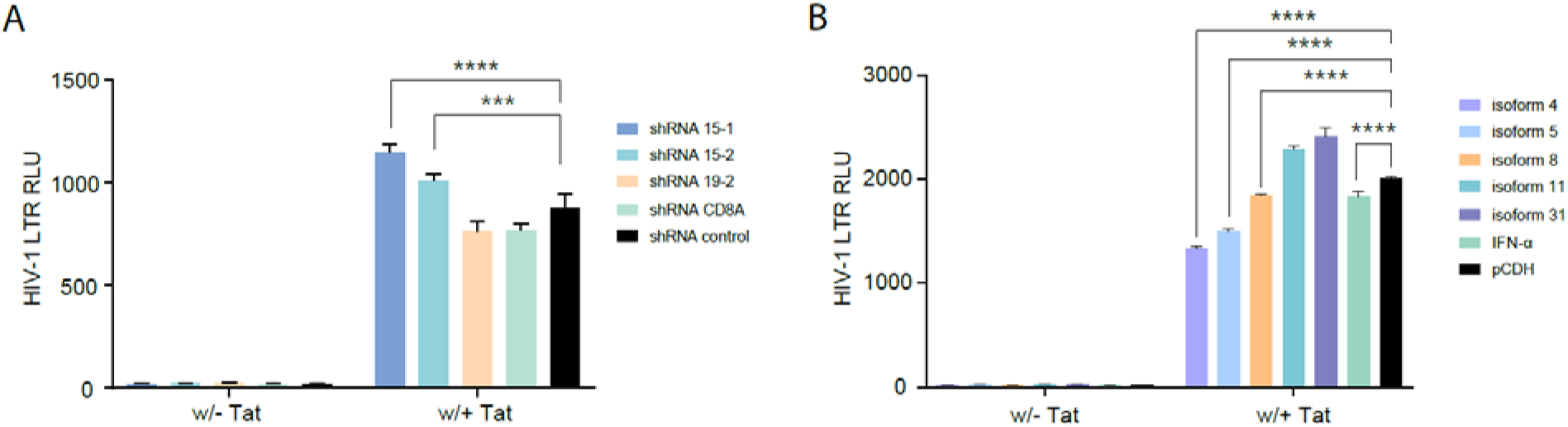
ICER regulates HIV-1 LTR promoter activity in a Tat-dependent manner. The activity of HIV LTR promoter was evaluated by LTR-driven luciferase reporter activity. (A) TZM-bl cells was stably transduced with shRNAs against ICER exon 15 (sh15-1, sh15-2) or 19 (sh19-2), or shRNAs against CD8A; the empty lentivirus was used as a vector control. (B) TZM-B1 cells was stably transduced with ICER isoform 4, 5, 8, 11, or 31; IFN-α was using as a positive control; the pCDH empty lentivirus was used as a vector control. These cells were transfected with pQCXIP-Tat vector or the empty one. Histograms show results in three independent experiments. Each bar represents mean ± SD. One-way ANOVA followed by Holm-Sidak’s multiple comparisons test. \*\*\**p* < 0.001, **** *p* < 0.0001.

## DISCUSSION

The main characteristics of ECs are undetectable viral load in blood and a steady CD4+ T cell count in the absence of ART. Understanding the mechanism for the natural control of HIV replication in ECs may yield new strategies for the treatment of HIV/AIDS. Here, we used a transcriptomic approach to compare gene expression in monocytes isolated from ECs and non-ECs, controlling for ART and HIV infection status. We found increased expression of specific, small isoforms of the CREM gene, all of which encoded ICERs.

To determine if the differentially expressed genes in ECs confer HIV resistance phenotypes to other cells, we overexpressed the most significantly altered genes. However, we failed to observe differences in HIV infection after overexpression of the differentially expressed genes. While individual overexpression of this gene subset did not influence infection in this model cell line, it is possible that depletion of these factors or investigation in a different model may identify unappreciated roles in replication. Alternatively, these genes may be representative of a gene expression program that renders cells resistant to infection, though these particular genes are not effectors in and of themselves. CREM is expressed in most immune cells, and encodes for many distinct protein isoforms that have broad impacts on multiple signaling pathways and organ functions. CREMα, for example, was found to perform critical functions as both epigenetic and transcriptional regulators for cytokine production in T lymphocytes [25]. Within the CREM family, the inducible cAMP early repressors (ICERs) are specific, short isoforms encoded in the latter half of the gene. ICERs inhibit T-cell activation, suppress proinflammatory cytokine production in macrophages [26, 27], control systemic autoimmunity in systemic lupus erythematosus (SLE) [28], and increase apoptosis via antiapoptotic protein expression [29].

Due to the CREM gene encoding many alternatively spliced transcript variants that are regulated posttranscriptionally [19], it is essential to understand CREM alternative splicing and isoform enrichment to interpret transcriptional changes. The IF and PSI calculations for the CREM gene splice sites showed that ECs widely use CREM exons 15, 16, and 18. Promoters located upstream of exons 15 and 16 drive specific ICER expression while exon 18 encodes a short 12-amino-acid segment enriched in acidic amino acids [20, 30]. The location of the promoter results in ICER proteins containing DNA-binding domains (DBDs) but lacking the upstream transactivation domain of CREM [20]. Additionally, the enriched ICERs in ECs contain intrinsically disordered regions (IDRs). IDR segments include a high proportion of polar or charged amino acids and lack a unique three-dimensional structure, permitting highly specific trigger signaling events while facilitating rapid dissociation when signaling is completed [31].

To determine the function of ICER in HIV infection, we knocked out the exons of ICER using HuT78 cell line. The knockout of ICER resulted in significantly increased viral replication and decreased cell counts after HIV-1 infection. However, knockout of CREM exon 13, which was wildly used in CREM isoforms, but absent in ICER isoforms, didn’t change the HIV-1 infection. We further verified the ICER function in primary CD4+ T cells. Knockdown ICER exons 15 and 19 resulted in increases in HIV-1 infection in primary CD4+ T cells across a range of HIV-1 strains, including both CCR5- and CXCR4-tropic viruses. Due to the short nucleotide fragment length of CREM exons 16 or 18 [20, 30], we could not identify effective shRNA against these exons. The ability of ICER to control HIV infection was also verified through overexpression of eight different ICER isoforms in primary CD4+ T cells using a lentiviral expression system. Although all ICER isoforms are highly expressed in HEK 293T cells, only some ICER isoforms were well-expressed in primary CD4+ T cells. Consistent with the knockdown results, the EC-specific ICER isoforms inhibited HIV-1 replication in primary CD4+ T cells.

These findings were further supported by small molecule agonists of ICER. Small molecule agonists can rapidly induce the ICER proteins, which subsequently serve as endogenous inhibitors of gene transcription by competing for binding with cAMP response element (CRE) sequences [32]. After treatment with ICER agonists PGE2 or 8-Br-cAMP, both ICER agonists strongly increased primary CD4+ T cell resistance to HIV infection. These data are consistent with a model in which alternative splicing at the CREM/ICER locus in ECs provides protection against infection and decreases viral setpoints. However, how this unique transcriptional profile is established, what effector genes are responsible for the observed protection, and what other genes may also contribute to this phenotype are yet to be understood.

In conclusion, increased ICER isoform RNA expression was identified in ECs. The results of knockout of ICER in HuT78 CD4+ T cell line, knockdown or overexpression of ICER in primary CD4+T cells, indicated that ICER is an HIV host factor that may contribute to HIV resistance of ECs. The alternative splicing of ICER for resistance to HIV is distinct from previously reported antiviral mechanisms. Understanding the role of ICER in HIV viral replication may help elucidate a novel target for treatment of HIV.

## MATERIALS AND METHODS

### Subjects

This study was conducted using cryopreserved peripheral blood mononuclear cells (PBMCs) from 10 ECs, 12 ART-treated HIV individuals, 11 ART-naïve HIV individuals, and 11 healthy individuals. Samples from ECs donors were received from National Institute of Allergy and Infectious Diseases (NIAID, from Dr. Stephen Migueles and Dr. Mark Connors). ECs had been infected by HIV more than 8 years with peripheral CD4+ T cell counts above 500 cells/μl and plasma HIV RNA less than < 50 copies/ml in the absence of ART. The clinical characteristics of HIV individuals are shown in Supplementary Table 1. Twelve samples from aviremic HIV-infected individuals on ART for at least 24 weeks (a single instance of ≤ 500 copies/ml was allowed) were collected from the Medical University of South Carolina (MUSC), clinical division of infectious diseases. Eleven HIV+ ART-naïve samples (plasma HIV RNA levels 1,000-600,000 copies/ml) were collected from Case Western Reserve University (from Dr. Michael Lederman) and the MUSC, clinical division of infectious diseases. Eleven healthy individuals were recruited from MUSC. The study received ethical approval from the MUSC review board. All recruited participants for this study provided written consent.

### RNA library preparation and sequencing

Human primary monocytes were obtained from PBMCs of 44 subjects. In brief, monocytes were enriched by negative selection using a Pan Monocyte Isolation Kit (Miltenyi Biotec, Bergisch Gladbach, Germany). Next, the isolated monocytes were further enriched by positive selection using CD14 MicroBeads (Miltenyi Biotec) according to the manufacturer’s instructions. The purity of the enriched monocytes was verified by flow cytometry to be greater than 95% for the next step. Total RNA was extracted from the purified monocytes with the RNeasy Plus Mini Kit (Qiagen, Hilden, Germany) according to the manufacturer’s protocol. The RNA purity was checked by a Nanodrop 2000 (Thermo Fisher), and the RNA integrity and size distribution were qualified by agarose gel electrophoresis and an Agilent 2100 Bioanalyzer (Agilent Technologies, Santa Clara, CA). Next, mRNA was enriched from total RNA by poly-T oligo-attached magnetic beads. The purified mRNA was fragmented randomly by the addition of fragmentation buffer. First-strand cDNA was generated using random hexamer primers and M-MuLV Reverse Transcriptase (RNase H). Second-strand DNA was synthesized using DNA polymerase I. The fragmented cDNA was purified using AMPure XP beads (Beckman Coulter, Brea, CA) and converted into blunt-ended DNA. The adaptors from the NEBNext Multiplex Oligos Kits (NEB, Ipswich, MA) were ligated to the adenylated 3’ ends of the DNA fragments. End-repaired DNA library fragments were again purified using the AMPure XP system. After several rounds of PCR amplification to enrich the cDNA fragments, the library was purified by AMPure XP beads. The purified library was quantified using a Qubit 2.0 fluorometer (Thermo Fisher), the average length was detected by an Agilent 2100, and the library concentration was quantified by real-time quantitative PCR (TaqMan Probe). The libraries were loaded onto a flow cell and subjected to paired-end sequencing on a HiSeq 2000 system (Illumina, San Diego, CA).

### Analysis of RNA sequencing data

RNA-sequencing data were processed and analyzed through a series of steps. First, Trimmomatic (v0.39) [33] was used to remove adaptors and trim low-quality bases. The quality of the processed reads was checked using FastQC before and after preprocessing. The reference genome GENCODE v29 (GRCh38.p12) and corresponding gene model annotation files were downloaded from GENCODE (gencodegenes.org). STAR (v2.7) [34] was performed using the default parameters to build the indexes of the reference genome and align the paired-end clean reads to the reference genome. Transcripts per kilobase million (TPM) were calculated based on the length of the gene and read count mapped to the gene. Differential expression between two groups (ECs vs. Healthy, ECs vs. ART, ECs vs. ART-naïve) was identified by the R Bioconductor package DESeq2 (v1.24.0) [35]. The resulting P-values were adjusted using Benjamini and Hochberg’s approach for controlling the FDR. Genes with an adjusted P-value < 0.05 were assigned as differentially expressed. The protein network analysis was performed via STRING (http://string-db.org/), and interaction scores > 0.3 were considered. Sashimi plots were applied for the creation of exon usage across experiments. The structures of the exons of human CREM gene and alternative splicing-generated isoforms were generated via GETxPortal (http://gtexportal.org).

### Alternative splicing analysis

The PSI score was used for alternative splicing quantification by psichomics (1.8.2) [36]. The alternative splicing event annotations were performed using hg29 genome assemblies from MISO, rMATS, SUPPA, and VAST-TOOLS. The PSI value was calculated as the inclusion/(inclusion + exclusion) reads, where inclusion and exclusion indicated the different splice events. Inclusion and exclusion events were obtained by extracting relevant junction reads from the output of STAR (v2.7) after splice-aware mapping. Salmon (v0.14) [37], which is based on “quasi-alignments” for mapping RNA-seq reads during isoform quantification, was performed. We also used RSEM (v1.3.1) [38] to implement iterations of expectation maximization algorithms to assign reads to the isoforms; the references were the GENECODE v29 genome and GTF file annotations. Calculations of IFs and prediction of functional consequences were performed with IsoformSwitchAnalyzeR [39] using standard parameters. We subsetted the CREM gene from the dataset, and the filter criteria were alpha = 0.05 and dIF cutoff = 0.01 in the isoformSwitchTestDEXSeq() function to test the CREM isoform switch. Protein domain prediction was performed using the Hmmer webserver (https://www.ebi.ac.uk/Tools/hmmer/search/phmmer); the result was incorporated in the switchAnalyzeRlist via the analyzePFAM() function, and the functional consequences of isoform switches were predicted by the analyzeSwitchConsequences() function.

### Cells

The HEK 293T cells used for the lentivirus production were provided by Dr. Eric Bartee (MUSC), and HEK 293T was maintained in complete DMEM plus 10% FBS. HeLa-derived TZM-bl cells expressing CD4, CXCR4, and CCR5 were obtained from the NIH AIDS Reagent Program (from Dr. John C. Kappes, and Dr. Xiaoyun Wu) [40].

### Lentiviral and retroviral vectors and molecular clones

The proviral infectious molecular clones included pNL4-3 (from Dr. Malcolm Martin) [41], pWT/BaL (from Dr. Bryan R. Cullen) [42], pNL (AD8) (from Dr. Eric O. Freed) [43], pYK-JRCSF (from Dr. Irvin SY Chen and Dr. Yoshio Koyanagi) [44], and p89.6 (from Ronald G. Collman) [45]. HIV-1 IIIB virus was provided by Dr. Robert Gallo [46]. All the above vectors and molecular clones were obtained from the NIH AIDS Reagent Program.

### Virus and virus-like particle production

Recombinant lentiviral particles, virus-like particles, and replication-competent HIV-1 were produced by transfection into 293T cells with the TransIT-Lenti transfection reagent (Mirus Bio, Madison, WI) at a ratio of 1:3 (DNA: TransIT-Lenti). 293T cells were seeded onto T25 cell culture flasks the day before transfection and grown to 80-95% confluence at the time of transfection. The overexpression vector (pCDH-puro controls, recombination vectors containing target genes) was cotransfected with pLP1, pLP2, and pLP/VSVG vectors at a ratio of 10:5:3:2. The virus with full replication capability was produced by transfection with a total of 5 μg of CXCR4/CCR5-tropic infectious molecular clone (pNL4-3, pNL (AD8), pWT/BaL, pYK-JRCSF, p89.6). After 18 h of transfection, the cell culture media were replaced with fresh media. Supernatants were harvested 48 h after transfection, passed through 0.45 μm syringe filters (BD), and treated with Benzonase (25 U/ml, EMD Millipore, Burlington, MA) for 1 h at 37°C to remove residual plasmid DNA. To generate HIV IIIB infection, primary CD4+ T cells were stimulated with PHA (4 μg/mL) in RPMI 1640 supplemented with 15% FBS and 20 IU/mL rIL-2 to promote virus production. Viral particle concentrations were determined by HIV-1 p24Gag ELISA (ZeptoMetrix, Buffalo, NY) or by determining the infectious titers in TZM-b1 cells. The viral stock was stored in aliquots at −80°C.

### Transduction of primary human CD4+ T cells with lentiviral vectors

Gene knockdown and overexpression were achieved using a lentivirus-mediated gene transfer system. For primary CD4+ T cells, PBMCs were isolated from whole blood of healthy human donors by Ficoll centrifugation. CD4+ T cells were isolated from PBMCs via the Easysep Human CD4+ T-cell negative enrichment kit (STEMCELL Technologies). The isolated human CD4+ T cells were cultured at 2.5 × 10^6^ cells/ml in complete RPMI medium (cRPMI), consisting of RPMI-1640 supplemented with 10% fetal bovine serum (FBS, vol/vol), 50 μg/ml penicillin/streptomycin, 1 mM sodium pyruvate, and 5 mM 4-(2-hydroxyethyl)-1-piperazineethanesulfonic acid (HEPES) in the presence of 20 U/ml IL-2. These cells were immediately stimulated with plate bounded anti-CD3 (UCHT1, coated overnight with 10μg/ml αCD3 (UCHT1, Tonbo Biosciences)) in the presence of 5 μg/ml soluble anti-CD28 (CD28.2, Tonbo Biosciences) at a final concentration of 2.5 × 10^6^ cells/ml. After stimulation for 3 days, cells were collected and replated into F-bottom, 96-well plate at 5 × 10^5^ cells/well in cRPMI with 20 U/ml IL-2. Next, 1 X 10^8^ reverse transcription units (RTUs) of lentivirus were added to the top. The plate was spun for 1 h at 1100 g, 32 °C, and then cultured at 37 °C in a 5% CO2 incubator. The cells were collected at the next day, washed, and replated into a 48-well plate. Cells were restimulated with anti-CD2/anti-CD3/anti-CD28 (T cell activation beads, Miltenyi Biotec) to achieve a 1:1 bead/cell ratio in accordance with the manufacturer’s instructions for cell expansion. Three days after transduction, puromycin (2 μg/ml) was added to allow puromycin-resistant cells to expand. After 3-day selection, the plate was kept at a 45 degree angle for 10 minutes; the aggregated cells were proliferated living cells settled at the bottom; then the aggregated living cells were slowly transferred into another 48 well plate. The cells count and viability were assessed using Muse Count & Viability Kit (Luminex Corporation, Austin, TX) and detected by Muse Cell Analyzer (MilliporeSigma, Burlington, MA). Next, the cells were transferred into U-bottom, 96-well plates at 1 X 10^5^ cells/well in cRPMI with 20 U/ml IL-2 and challenged with replication competent HIV-1.

### Infection with retrovirus virus

We placed 1×10^5^ primary CD4+ T cells in each well in a 96-U-well plate and inoculated them with HIV NL4-3 suspension. After inoculating CD4+ T cells with the AD8, JR-CSF, 89.6, BaL, or IIIB virus, the plate was spun for 1 h at 800 g, 32 °C, and then cultured at 37 °C in an incubator. After transferring the cells to a 96-U-well plate, the cells and HIV were incubated overnight at 37°C. The infected cells were washed with 200 μl of fresh culture medium. On day 3 or 5, the cells were stained with LIVE/DEAD Aqua blue (Thermo Fisher) for 20 min at 4°C to detect and exclude dead cells. Next, the cells were fixed, permeabilized and stained for intracellular p24 using anti-p24-PE antibodies (Beckman Coulter). Data were acquired using a FACSVers flow cytometer (BD Biosciences, San Jose, CA) and analyzed using FlowJo software 10.0.6.

### CREM/ICER western blot

The primary CD4+ T cells were lysed in the RIPA buffer with protease/phosphatase inhibitor cocktail (Cell Signaling Technology, Danvers, MA). After determining the protein concentration for each cell lysate, an equal volume of 2X Laemmli sample buffer with β-mercaptoethanol was added to equal amounts of cell lysate. After denaturation at 95°C for 5 min, the cell lysate in the sample buffer was loaded with equal amounts of protein into the wells of an Any kD mini-PROTEAN protein gels (Bio-Rad, Hercules, CA). Next, the proteins were transferred from the gel to a 0.2 μm Immobilon-PSQ PVDF Membrane (Millipore). The membrane was blocked for 1 h at room temperature and incubated overnight at 4°C with CREM/ICER antibody (clone 3B5) (LifeSpan BioSciences, Seattle, WA). Mouse anti-human β-actin was used as the reference. After washing the membrane three times using TBST, an anti-mouse HRP-conjugated secondary antibody was incubated for an additional hour. The HRP activity was determined using the SuperSignal West Femto Maximum Sensitivity Substrate (Thermo Fisher) and scanned the blot using a ChemiDoc MP Imaging System (Bio-Rad).

### Generation of stably transduced cell lines using CRISPR/Cas9

HuT78 cells were transfected with 2.5 μg of pSpCas9(BB)-2A-GFP sgRNA expression vectors (a gift from Feng Zhang, Addgene plasmid # 48138) [47] using Lipofectamine 3000 (Thermo Fisher, Waltham, MA) and incubated at 37°C with 5% CO2. The transfection procedure was performed following the manufacturer’s instructions. After transfection for 48 h, single GFP-positive cells were sorted into 96-well U-plates using a BD FACS Ari II flow cytometer and expanded for two weeks. The individual clones were screened by sequencing. For each exon knockout, we selected clones that had frameshift (3n+1 bp) mutations in both alleles. The clones had deleterious frameshift mutations in both alleles, but lacked large deletions or any change to the intronic sequence.

### Statistical analysis

Conventional measurements of central location and dispersion were used to describe the data. Non-parametric Mann-Whitney’s U tests were applied to compare differences in continuous measurements between the two groups. A one-way ANOVA followed by Holm-Sidak’s multiple comparisons test was used to compare differences among more than two categorical groups. Significance for comparisons of individual gene expression in RNA-seq data was tested using the Wald test in DESeq2 package; *p* values denoted were adjusted using Benjamini-Hochberg correction. Comparison analysis was performed using R (version 3.3.1) or GraphPad Prism 8. *p* ≤ 0.05 was considered statistically significant.

## Acknowledgment

We acknowledge Dr. Michael Lederman from Case Western Reserve University for providing human specimens from ART-naïve HIV-infected individuals as well as reviewing this manuscript. We acknowledge Dr. Mark Connors from National Institute of Allergy and Infectious Diseases for providing human specimens from HIV-infected elite controllers.

## Funding

National Institute of Allergy and Infectious Diseases grant AI128864 (W. J.)

South Carolina Clinical & Translational Research Institute with an academic home at the Medical University of South Carolina CTSA NIH/NCATS grant UL1TR001450 (W. J.)

Gilead Sciences Research Scholars Program in HIV (J.F.H.)

NIH grant K22 AI136691 (J.F.H.)

NIH grant R01 AI150998 (J.F.H.)

NIH-supported Third Coast CFAR P30 AI117943 (J.F.H.)

NIH-sponsored HARC Center P50 GM082250 (J.F.H.)

## Author contributions

Z.L. and W.J. conceived the study. Z.L. wrote the manuscript. Z.L., T.L., and W.C. performed experiments. Z.L. and M.L. analyzed data. M.L., ZY.L., Z.Y., J.F.H., J.Z., L.Y., and J.Z. conceived the study and revised the manuscript. S.M. provided the key human specimens and revised the manuscript.

## Data availability

RNA-seq data in this study have been deposited in the National Center for Biotechnology Information Gene Expression Omnibus (GEO) database under accession no. GSE157198.

## Supplementary Figures

**Figure S1.**
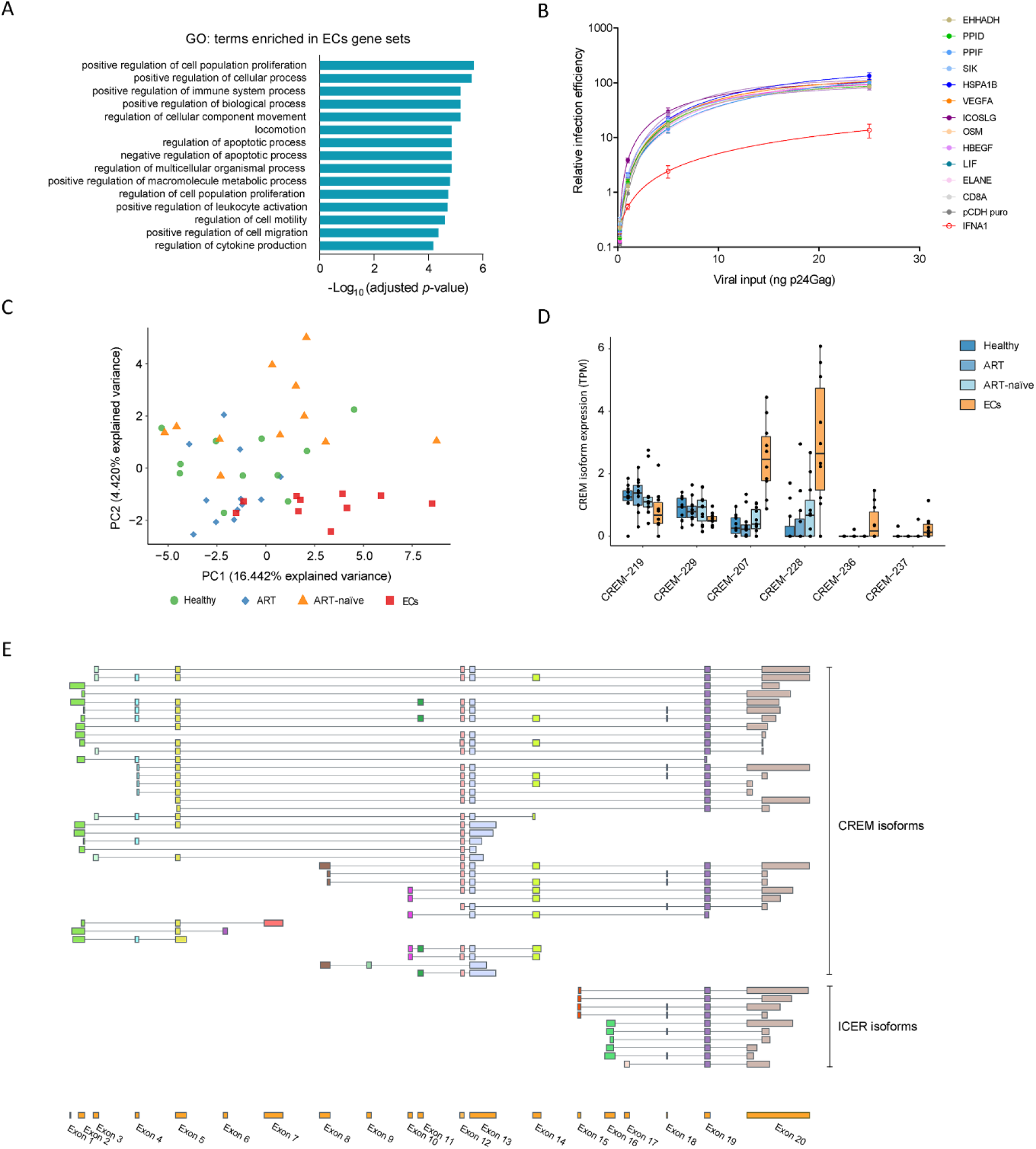
The differential gene expression profiles and alternative splicing features of the CREM/ICER gene. (A) Clustering of Gene Ontology (GO) termed function and pathways of the differentially expressed genes in ECs compared with the other three groups. (B) Candidate genes were screened in HuT78 CXCR4 cells. HuT78 CXCR4 cells were transduced with recombinant lentiviruses expressing the different candidate genes, CD8 (a negative control), and IFNA1 (a positive control) cDNAs; cells were selected using puromycin. HuT78 cells were infected with increasing viral inputs of HIV NL4-3, and infection efficiency was monitored by measuring Gag+ cells at 72 h after infection. Mean relative infection efficiencies with standard deviations from five replications are shown. (C) PCA was conducted to show alternative splicing events of CREM gene in ECs compared with the other three groups. (D) The alternative splicing of CREM/ICER gene was quantified by comparing the six predicted CREM/ICER isoforms using RNA expression (TPM). (E) Structure of the exons of human CREM/ICER gene.

**Figure S2.**
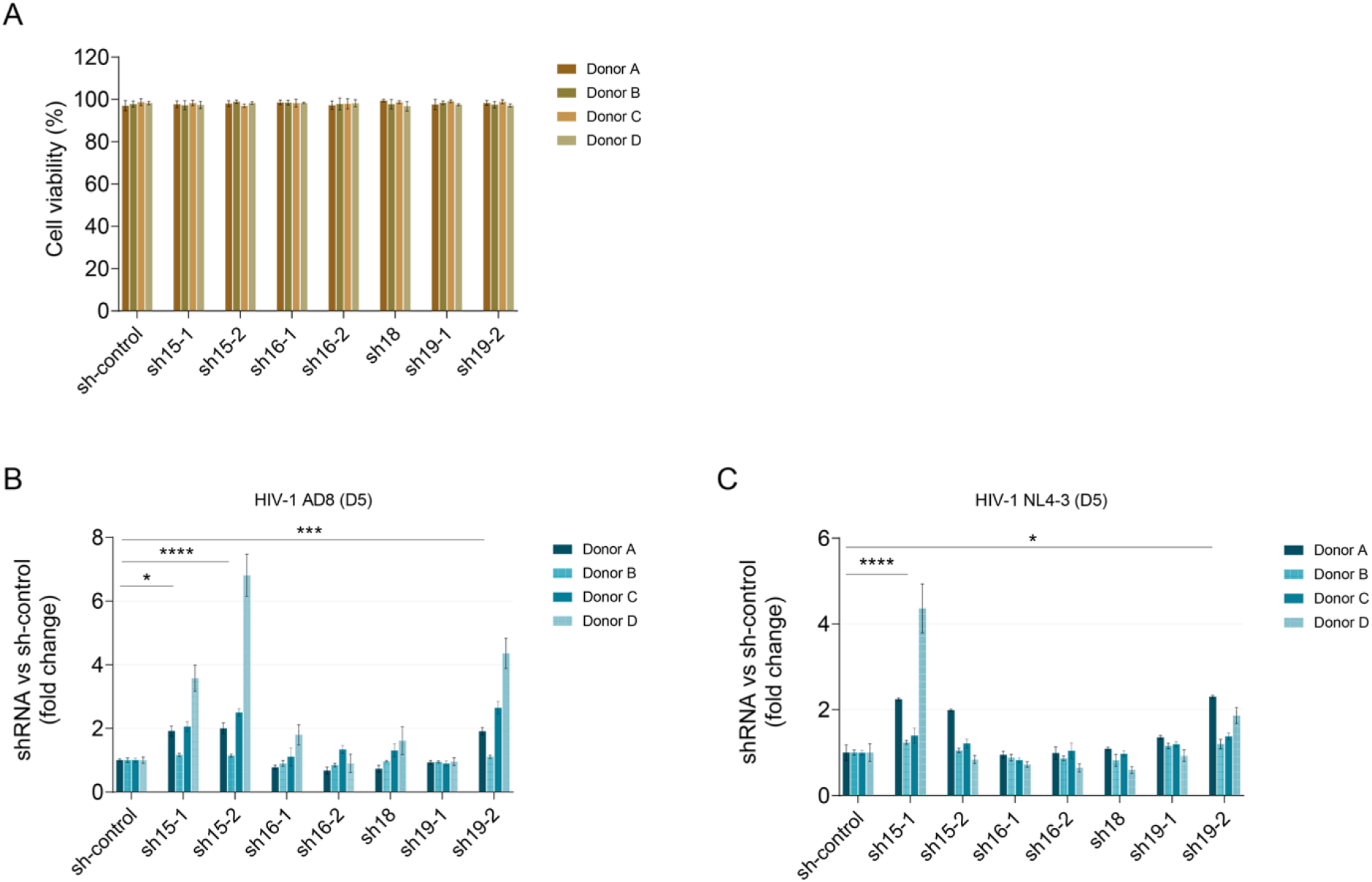
Increased HIV infection in primary human CD4+ T cells upon ICER knockdown. The ICER controlled both HIV AD8 and NL4-3 infection and replication. Primary human CD4+ T cells were activated using anti-CD2/CD3/CD28, transduced with ICER shRNAs against exons 15 (sh15-1, sh15-2), 16 (sh16-1, sh16-2), 18 (sh18), or 19 (sh19-1, sh19-2); empty lentivirus was used as a control (sh-control). The transduced cells were selected using puromycin. (A) The percentage of cell viability in primary CD4+ T cells transduced with ICER shRNAs or empty lentivirus control. Cell viability was determined using Muse Count & Viability Kit. (B, C) The primary CD4+ T cells were infected with HIV-1 AD8 (B) or NL4-3 (C) after knockdown of ICER exon 15, 16, 18, or 19 using shRNA. The effect of ICER shRNA on HIV infection was monitored by p24 staining on day 5 after viral challenges. Results are displayed as the fold changes of HIV-1 infection in ICER shRNA transduced T cells compared to those of sh-control transduced T cells. Histograms show results from four different donors. Each bar represents mean ± SD of triplicates for each donor. One-way ANOVA followed by Holm-Sidak’s multiple comparisons test. * *p* < 0.05, \*\*\**p* < 0.001, **** *p* < 0.0001.

**Figure S3.**
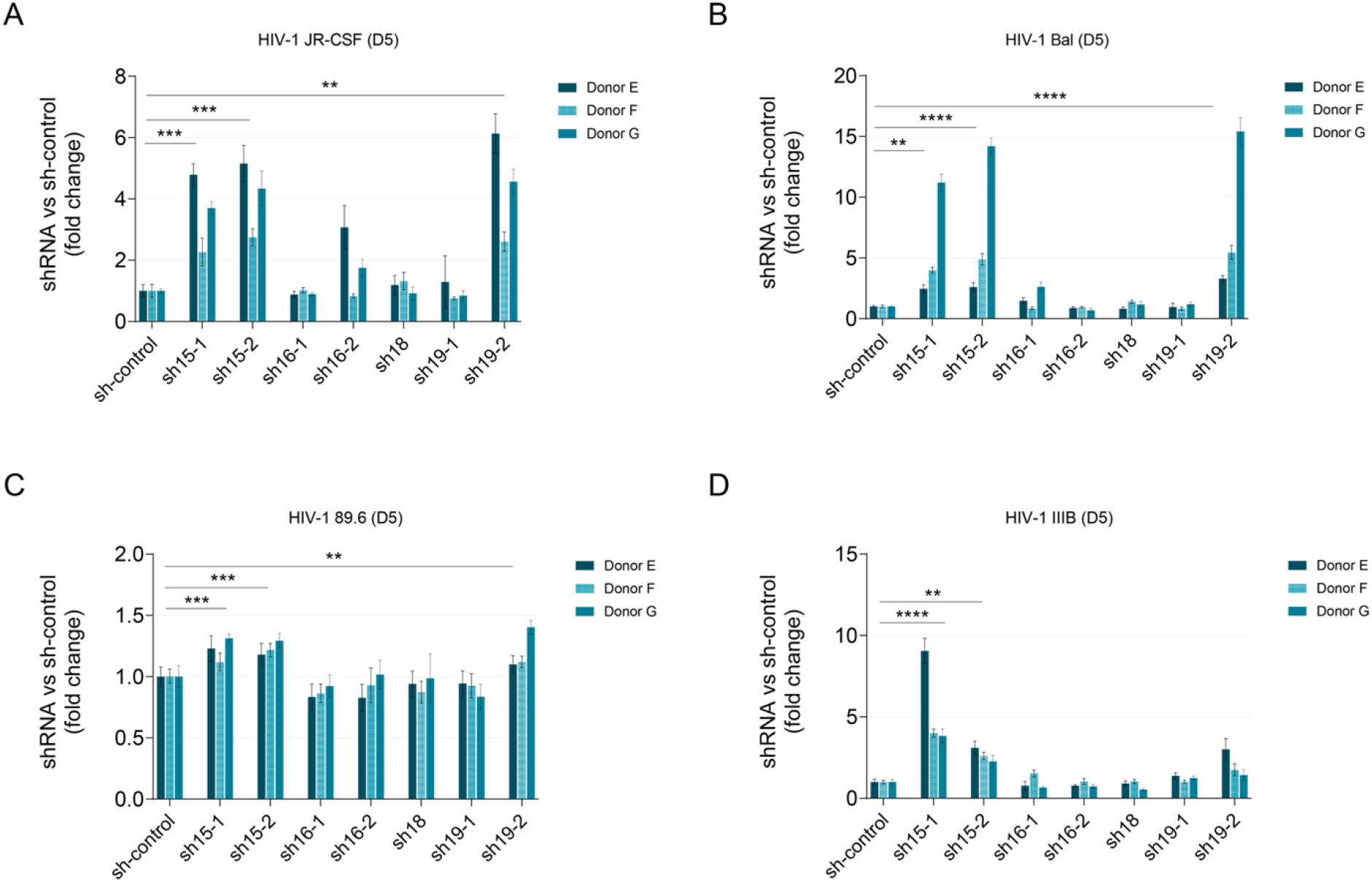
ICER controls different strains of HIV-1 infection in human primary CD4+ T cells. Primary CD4+ T cells were activated using anti-CD2/CD3/CD28, transduced with ICER shRNAs against exons 15 (sh15-1, sh15-2), 16 (sh16-1, sh16-2), 18 (sh18), or 19 (sh19-1, sh19-2); empty lentivirus was used as a control (sh-control). Cells were selected by puromycin. The transduced cells were then infected with HIV-1 JR-CSF (A), BaL (B), 89.6 (C), and IIIB (D). The viral replication efficiency was evaluated by p24 staining on day 5 after HIV infection. Results are displayed as the fold changes of HIV-1 infection in ICER shRNA transduced T cells compared to those of sh-control transduced T cells. Histograms show results from three different blood donors. Each bar represents mean ± SD of triplicates for each donor. One-way ANOVA followed by Holm-Sidak’s multiple comparisons test. ** *p* < 0.01, *** *p* < 0.001, **** *p* < 0.0001.

**Figure S4.**
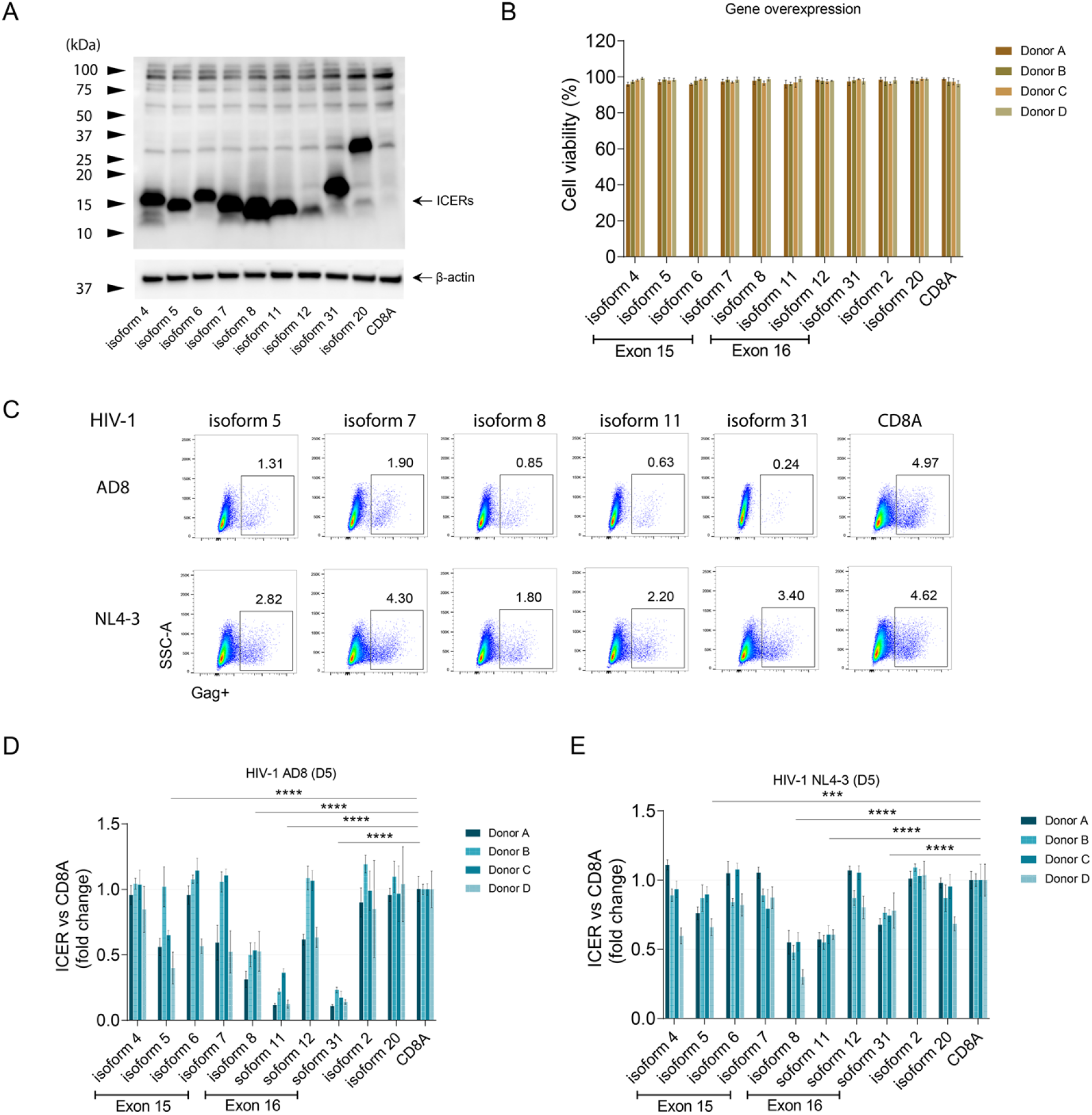
Resistance to HIV infection in human primary CD4+ T cells after ICER isoform overexpression. Primary CD4+ T cells were activated using anti-CD2/CD3/CD28, transduced with CREM/ICER isoforms using lentivirus system, and selected by puromycin. (A) The transfected HEK 293T cells with overexpression of CREM/ICER isoforms or with the empty plasmid. (B) Histograms represent the percentage of cell viability in primary CD4+ T cells overexpressed CREM/ICER isoforms. (C) Dot plots representing percents of Gag+ cells on day 3 after HIV AD8 or HIV NL4-3 infection upon overexpression of CREM/ICER isoforms or control gene in primary CD4 T cells. (D, E) The transduced primary CD4+ T cells were infected with HIV-1 AD8 (D) and NL4-3 (E) for 5 days. Results are displayed as the fold changes of HIV-1 infection in CREM/ICER transduced T cells compared to those of CD8A gene transduced T cells. Histograms show results from four different blood donors. Each bar represents mean ± SD of triplicates for each donor. One-way ANOVA followed by Holm-Sidak’s multiple comparisons test. * *p* < 0.05, \*\**p* < 0.01, *** *p* < 0.001, **** P < 0.0001.

**Figure S5.**
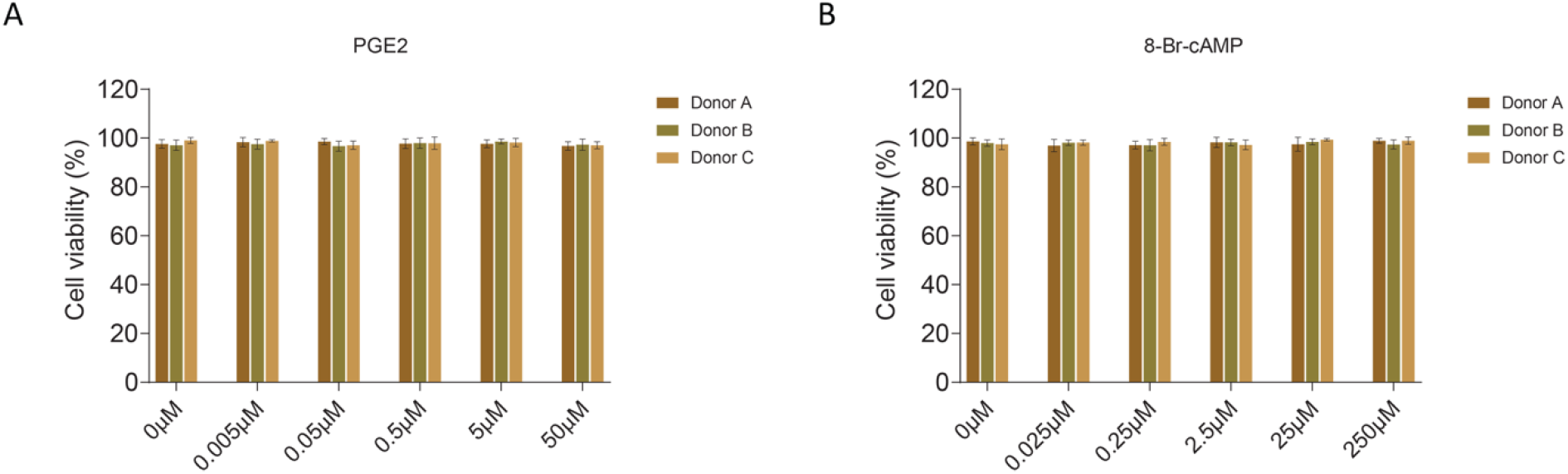
The cell viability of primary CD4+ T cells after treatment with ICER agonists. Anti-CD2/CD3/CD28 activated primary CD4+ T cells were treated with PGE2 (50 μM-0.005 μM) (A) or 8-Br-cAMP (250 μM-0.025 μM) (B) for 16-24 h, and the cell viability was shown. Histograms show results from three different blood donors. Each bar represents mean ± SD of triplicates for each donor.

